# The impact of coinfection on population stability and chaos

**DOI:** 10.64898/2026.03.06.710155

**Authors:** Fernando Melara Barahona, Edith Simpson, Ann T. Tate

**Affiliations:** Department of Biological Sciences, Vanderbilt University, Nashville, TN, USA; Evolutionary Studies Initiative, Vanderbilt University, Nashville, TN, USA

**Keywords:** LPA model, flour beetles, parasite facilitation, antagonism, discrete time model

## Abstract

Parasites play an outsized role in mediating the persistence and stability of host populations. Flour beetles (*Tribolium* spp.) have long served as classic examples of population dynamics under both disease-free and infected conditions, with elegant combinations of theory and experiments demonstrating, for example, that cannibalism rates can push populations from stability to chaos. As with most organisms in nature, however, flour beetles rarely face just one parasite species, and co-infecting parasites can antagonize or facilitate each other through resources and immunity. To test the prediction that non-neutral interactions would qualitatively alter population stability, we first raised flour beetles (*Tribolium castaneum*) in infection-free, single-infection, or coinfection microcosms and quantified relative prevalence and parasite intensity. Next, we reworked a classic stage-structured discrete-time model to include single and multiple infections and performed sensitivity and bifurcation analyses to identify the most important (co)infection-associated parameters for population stability. The model predicts that stability is highly sensitive to parasite transmission mode regardless of infection multiplicity, but facilitation among parasites rapidly drives populations into oscillations and chaos under realistic conditions. This study identifies an important mechanism for explaining population variability and highlights the importance of within-host mechanisms for driving dynamics at higher levels of biological organization.

## Introduction

Parasites are ubiquitous in nature and are central to the regulation of organismal populations. Because they steal resources from host reproduction and promote disease-induced mortality, among other demographically relevant effects, they can alter the stability of populations [1,2] impose apparent competition among host species [3], and even drive local populations to extinction [4]. A classic proof of principle stems from an experiment that eliminated strongyle nematodes from red grouse populations, promoting population stability and thereby demonstrating the influence of the parasite on the promotion of host population cycles [5].

While a single parasite species can certainly impose dramatic ecological effects on its host population, hosts rarely harbor just one parasite infection over the course of their lifespan [6]. After all, most populations encounter an assemblage of specialist and generalist parasites, resulting in multiple infections or even coinfection by multiple parasites at the same time [7]. Within the host, parasites can facilitate one another through immunosuppression or antagonize each other through competition for resources [8,9] or cross-immunity [10]. These interactions, in turn, affect the fitness, ecological dynamics, and force of infection of individual parasite species. A theoretical model, for example, predicted that facilitation among generalizable microbial strains could increase the probability of secondary pathogen invasion at the population scale [11]. At the same time, within-host parasite interactions alter parameters relevant to host population dynamics, such as disease-induced mortality [12].

Alongside interaction type, transmission mode must also be considered when interpreting epidemiological outcomes at a population scale [3,13,14] because it is an essential part of the parasite life cycle and can interact in important ways with host mortality and other epidemiological parameters [15,16]. Obligate killer pathogens, for instance, cause acute infections that require host death for successful transmission. Such is the case of *Bacillus thuringiensis* (Bt), an entomopathogen known to control invertebrate populations via the production of lethal toxins [17,18]. Directly transmitted parasites, on the other hand, need to ensure host survival to maximize their fitness [19–22]. Eugregarines are a group of directly transmitted protozoan parasites, ubiquitous in insect populations, that typically impose little pathology beyond resource exploitation; this allows them to find a mate and reproduce all while living comfortably in the gut of their living host [23]. Given these opposing transmission imperatives, the outcome of co-infections for both hosts and parasites can be particularly difficult to predict for mixed transmission mode assemblages.

Thus, co-infection introduces a degree of entropic variability that could lead to changes in the stability of population dynamics in the focal species, depending on the combination of parasites involved and the net effect of facilitation and antagonism on host and parasite population parameters. However, it remains unclear whether multiple parasite infections simply reinforce shifts in population stability compared to single infections, or if co-infected populations experience a fundamentally different stability landscape. This gap in knowledge is difficult to address because it requires data from a system where infection, host demographic processes, and stability can be measured directly. While many theoretical frameworks incorporate within-host processes and their consequences for transmission and disease prevalence [24–28], few take a broader view of their impact on overall population stability across a generalizable parameter space. A suitable model must recapitulate the basic life-history traits, stage-specific demographic processes, density-dependent regulation, transmission mode, ecological interactions, and infection structure of the focal model system – all while remaining tractable for proper analysis.

Flour beetles of the genus *Tribolium* have long been a cornerstone model in population biology and stability analysis due to their well-documented life histories and ease of experimental study [29]. Their tractability for infection, culturing, and censusing, combined with measurable density dependence and a relatively short life cycle, make them ideal for the integration of empirical studies and theory [30–32]. As a result, robust mathematical models have been developed to explore population fluctuations and stability, including the larva-pupa-adult framework (LPA) [33]. This discrete-time model divides populations by developmental stages, linking demographic processes with nonlinear population feedbacks. Parameterized with experimental data, the model demonstrates that long-term population dynamics hinge on nonlinear interactions between compartments, with cannibalism at the core of this process. By regulating recruitment, cannibalism directly influences population size. Strengthening this feedback loop enhances stability, while altering it introduces oscillations, chaos, and delayed population growth [34]. In essence, cannibalism acts as a regulatory force, determining the system’s eventual state [14,34]. Subsequent theoretical studies have expanded this work to include spatial extensions, global dynamical analyses, and modern computational approaches [14,31,32]. These studies collectively highlight the intricacy and broad applicability of the model.

Co-infection perturbs the same parameters – survival, reproduction, and effective density – that cannibalism stabilizes [35–37], suggesting that even small infection-induced perturbations could force the system toward new attractors [31]. Recent experimental work in *Tribolium* provides proof of principle that interactions between Bt bacteria and eugregarines can modify host development, mortality, and body condition under co-infection [38], linking within-host processes to traits that influence transmission and demographic performance and introducing potentially destabilizing forces that differ from the effect of single infections [39–41].

In this study, we incorporated single- and co-infection dynamics into the LPA model to examine how infections shape system stability. We first performed experiments in *T. castaneum* infected or co-infected with an obligate killer (the protozoan *Farinocytis tribolii*) and the directly transmitted eugregarines to quantify any interaction effects among parasites on prevalence and intensity. We then turned to the model to evaluate stability profiles under disease-free, single infection, and co-infection scenarios. We started by bifurcating over adult-on-pupa cannibalism rate, a key parameter modifying stability in disease-free conditions [33]. We then systematically evaluated the roles of transmission mode and parasite interaction modes (antagonism vs. facilitation) on bifurcation structure and routes to chaos. Overall, our results demonstrate that co-infection stability profiles differ in important ways from single infection scenarios, emphasizing the need to account for both transmission mode and the mechanisms of within-host interactions when predicting the impact of multiple infections on host population dynamics.

## Methods

### Preparation of experimental infection mesocosms

Previous experiments demonstrated an increase in disease-induced mortality from infection with an obligate killer (Bt) when *Tribolium* beetles were coinfected with eugregarines [38], but it was unclear whether this interaction facilitated or stymied parasite transmission and prevalence, not in the least because *Tribolium* are relatively resistant to the Bt cry toxins that are required for sustained transmission. Therefore, we turned to a different obligate killer with a dependable natural transmission route (*F. tribolii*) to determine the prevalence of eugregarine and *Farinocystis* parasites in *T. castaneum* under single and co-infection conditions in population mesocosms. 100 uninfected adult beetles from the lab-reared *Snave* population were transferred to a mating jar containing 150 grams of whole wheat flour mixed with 5% w/w brewer’s yeast and allowed to produce eggs for a 24 to 48-hour period. From this common pool of eggs we distributed 50 eggs per experimental block (n = 3) into one of four treatment groups: 1) a control group exposed to uninfected flour, 2) a group exposed to flour containing eugregarine oocysts, 3), a group exposed to flour containing *Farinocystis* spores, and 4) a coinfected group in which the beetles were exposed to both parasites. We prepared infected flour using a mortar and pestle to crush larvae previously infected with neogregarine parasites and release well-mixed oocysts into the flour. For eugregarines, infected flour was obtained by allowing infected larvae and adults to shed parasite-containing frass into the flour. Infected groups received 8g of infected flour with frass plus 4g of clean flour. Control groups received 8g of clean flour with frass (made by crushing clean larvae in clean flour) plus 4g of clean flour. All groups received a total of 12g of flour.

Larvae were collected approximately 14 days after exposure to assay for prevalence and eugregarine intensity. The gut and fat bodies of larvae were dissected in phosphate-buffered saline (PBS), stained with iodine, and inspected via microscopy [42]. Parasite prevalence was measured based on the presence or absence of one or more parasite stages within the host. Intensity was quantified as the number of eugregarine parasites in the beetle gut (0-100 were counted precisely but loads >100 were binned as such); *Farinocystis* intensity could not be reliably obtained due to the difficulty of quantifying non-oocyst stages.

### Statistical analysis

Analyses of experimental data were conducted in R version 4.3.2 [43]. Parasite prevalence was initially analyzed using generalized linear mixed models with a binomial error distribution and a logit link function, with parasite type included as the explanatory variable and block as a random effect. Since the effect of block was not significant, however, we dropped the mixed model and performed simple Chi Square tests on the combined data to compare prevalences across mesocosms. Even though we suspect that eugregarines follow a negative binomial distribution, we had too few positive samples in one of the mesocosms to get an accurate estimate of overdispersion so we relied on a Kruskal-Wallis test to compare gregarine intensity within positive samples across treatment groups. Statistical significance was assessed at α = 0.05 for all tests.

### Modifying the LPA model

We developed discrete time, stage-structured nonlinear infection models capturing life-history processes, density-dependent regulation, and environmentally mediated infection in *Tribolium castaneum*. Infection dynamics are inspired by the directly transmitted eugregarine protozoa that reside in the gut of flour beetles and other insects [38] as well as obligate killers like *Adelina tribolii*, the coccidian parasite from a classic demonstration of parasite-mediated apparent competition in this system [44], neogregarine protozoa (*Farinocystis tribolii*), and Bt, all of which form spores in the host hemocoel and require host death to achieve transmission. These common *Tribolium* parasites all spread via environmental stages and disproportionately affect the larval stage [22,42]. To investigate the impact of co-infection and transmission modes on host population stability, we build on the classic LPA framework, a stage-structured discrete time model that originally demonstrated the sensitivity of *Tribolium* population stability to cannibalism rates [33]. The LPA model describes the life cycle of beetles using three stochastic equations (Eqs. 1-3):

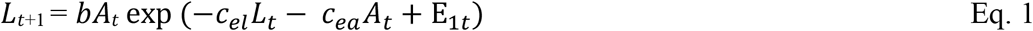

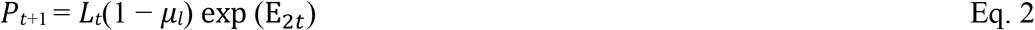

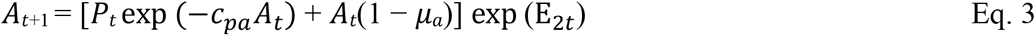

The model assumes a multivariate distribution with stochastic coefficients set to zero and uses a time step of two weeks, approximating the duration of larval and pupal beetle stages [33]. The first equation describes the dynamics of larvae at the next time step as a function of adult fecundity (*b*), the number of adults at time t, and the probability that eggs survive cannibalism by larvae (*c*_*el*_) and adults (*c*_*ea*_). Eq. 2 describes the number of pupae as a function of larval survival from the previous time step. The third equation tracks the number of adults after pupal eclosion, which is reduced by pupal cannibalism by adults (*c*_*pa*_) as well as the background mortality rate of existing adults (μ_a_).

Building on this framework, we retained the same three life-stage compartments but introduced explicit larval infection and parasite environmental compartments; since larvae are the main drivers of transmission and mortality for these parasites, we made the simplifying assumption that pupae and adults could not get infected. We assume that infection events occur randomly and independently among larvae, with a constant mean exposure rate over each time step, reflecting the use of exponential transmission in our model [45].

### Incorporating infection and transmission modes into the LPA model

To generalize the equations, we use index *i* to refer to the parasites in single-infection scenarios and index *j* to refer to the second parasite when both parasites are present in the population. L_S_ refers to uninfected larvae. The total larval population includes uninfected, singly infected, and coinfected subpopulations and is given by the equation:

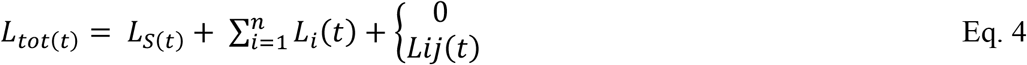

The number of new larvae at time t+1 (hereafter noted as *R*_*t*_) is identical to Eq. 1 except L_tot(t)_ replaces the L_(t)_ in Eq. 1. The proportion of larvae that are allocated to infected or uninfected compartments is determined by the infectiousness *β* of environmental spores (E). We assume that the probability of an encounter between just-hatched larvae and pathogens follows a Poisson distribution, where the probability of remaining uninfected is *e*^−*βiEi*^ [45]:

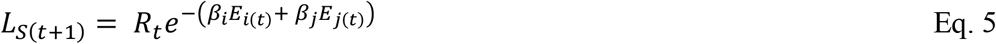

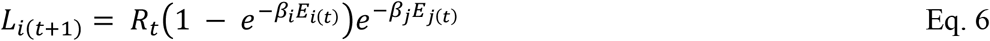

We represent the environmental stage spores with background decay rates *δ* and carrying capacities K_i_ through two equations that are differentially affected by larval disease-induced mortality rates (*α*_*i*_); larvae must be alive at the end of the time step to release directly transmitted spores (D), whereas larvae must die to release obligate killer (OB) spores:

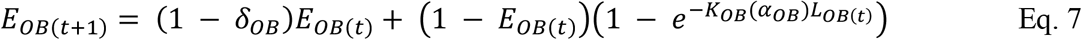

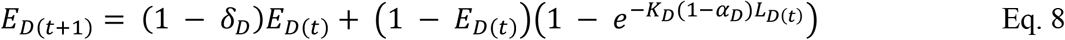

Co-infecting parasites can facilitate or antagonize each other via competition for resources, cross-immunity, or immunosuppression, among other mechanisms [3], ultimately affecting the production of transmission stage spores. To account for these interactions, we added an interaction coefficient *ϕ*to the exponential term of the environmental spore equations. A value less than one would indicate antagonism while a value greater than one would indicate facilitation; a value of one would indicate no (neutral) interaction among parasites. For example, the number of new environmental spores for a directly transmitted parasite *i* that is co-infecting a host with parasite *j* (regardless *j*’s transmission mode) would be represented as:

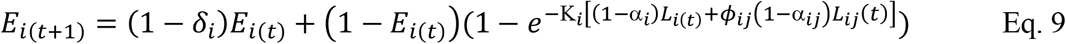

The full set of equations for each coinfection scenario explored in this study can be found in the Supplementary Materials.

The number of adults at t+1 is identical to Eq. 3, while the number of pupae is similar to Eq. 2 except for the loss of individuals to disease-induced mortality during the transition from infected larvae to pupae:

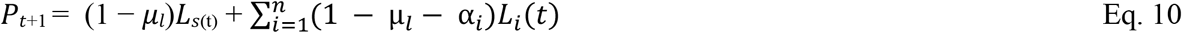

To prevent unrealistic environmental contamination values and account for over- and underflow, numerical clamps were anchored between 0 and 1.

### Forward and backward bifurcations

For these analyses, all simulations were performed in MATLAB v.R2025b [46]. Each time-step in this discrete-time model corresponds to one two-week period. We ran simulations for 3000-time steps to ensure robustness and convergence to long-term dynamics while discarding the first 200 time-steps to allow for stabilization. At each time-step, population states were updated in a fixed sequence. We conducted parameter sweeps at a relative frequency magnitude of 10^−8^, using (for example) 2000 evenly spaced points between 0 and 1 to represent cannibalism of pupae by adults (*c*_*pa*_). Each infection scenario included six multi-state systems: No infection, single infection with either a directly transmitted or obligate killer parasite, and co-infection with factorial combinations of directly transmitted and obligate killer parasites.

We first ensured that we could recreate the original *c*_*pa*_ bifurcation plots from [33] under disease-free conditions (all infection parameters set to zero) and used the same initial conditions and baseline parameter values for the infection scenarios unless otherwise noted in **Table S1**. Most parameter values were primarily derived from empirical data, and when estimates were necessary, a sensitivity analysis was conducted to ensure alignment with experimental prevalence and assess the impact of these estimates on model results.

Demographic parameters were selected to span biologically relevant ranges known to generate equilibria, periodic and quasiperiodic cycles, and chaotic dynamics [34]. For each parameter set, the model was iterated deterministically from the specified initial conditions for the full simulation duration. State variables were recorded at each time step. Infection prevalence for relevant scenarios was calculated as fraction of infected larvae within the total larval population. Time-series plots of prevalence were then used to assess whether infections persisted, died out, or exhibited transient dynamics, with a low prevalence threshold (1%) used as a reference for infection persistence. Differential stability across infection states was quantified as the difference between the maximum Lyapunov exponents of the co-infected and single-infection models, computed using the forward sweep.

### Largest Lyapunov exponent calculation

Using the same time-steps as in the forward and backward bifurcation plots, maximum Lyapunov exponents (LE) were calculated with a Benettin two-trajectory method [47]. Post-transient LE values were computed after discarding an initial transient period and subsequently plotted as a function of *c*_*pa*_, the sweeping control parameter to which the stability of disease-free dynamics is so sensitive [33]. A small perturbation in the order of 10^−8^ was applied to the state vector and the divergence was tracked over time. At each step, the perturbation was normalized, and the Lyapunov exponent was computed as the time-averaged exponential divergence rate.

### Largest Lyapunov exponent analysis in two parameter space

Maximum Lyapunov exponents were calculated using the same numerical pipeline as in our multi-infection state analysis. In this case, however, the analysis was extended to a two-parameter sweep to construct a Lyapunov exponent map. For each pair of control parameters, the system was run for 1000 time-steps, of which 100 were discarded as transient dynamics. A grid of 2000 evenly spaced values spanning the interval [0,1] was generated for each parameter, and the resulting Lyapunov exponent values were reassembled into a two-dimensional parameter space. The resulting LE values were organized and color coded using a continuous color scale, highlighting transitions between stable and unstable dynamical regimes. For each condition, parameter contributions to variability were quantified using the Sobol total-effect sensitivity index (*S*_*T*_) using MATLAB [46].

### Global sensitivity analysis

We used a variance based-Sobol framework with Saltelli sampling [48] to conduct a global sensitivity analysis and determine the parameters that most strongly influenced the largest Lyapunov exponent. Parameters were constrained to biologically plausible ranges (**Table S2**), and for each parameter set, the model was run forward to remove transient dynamics before estimating the maximum Lyapunov exponent. First-order Sobol indices assess the effect of changing each parameter on its own, while total-order indices capture the overall impact of each parameter, accounting for multi-order parameter interactions [48]. Uncertainty in these indices was estimated using bootstrap resampling to estimate confidence intervals.

### Stability analysis on the parasite interaction parameter

To further evaluate how facilitative and antagonistic interactions among two parasites influence population stability, we conducted shortened simulations (1000 time-steps), excluding the first 200 as transient dynamics. We performed forward parameter sweeps over the antagonism/facilitation coefficient (*ϕ*), assuming symmetry between interactions. A more robust functional form of the model, introducing *ϕ*as a multiplicative modifier on the contribution of co-infection to the infectiousness of spores in Eq. 5 and 6 (*ϕ*_*ij*_*β*_*i*_*E*_*i (t*)_), was included. We allowed parasite-specific contributions (*ϕ*_1_, *ϕ*_2_), defining neutrality as conditions under which co-infection generated no additional transmission benefit beyond baseline expectations. Extending this approach to explore the role of interaction structure, we also considered a symmetric interaction in which co-infection effects were represented by a single shared interaction parameter rather than parasite-specific terms. In this framework, *ϕ*= 1 represented neutral interactions. All parameter values were taken from **Table S1**, which provides complete parameter description, biological justification, empirical values, and parameter estimates.

## Results

### Empirical estimation of infection prevalence and eugregarine intensity under single and co-infection conditions

We dissected the gut and fat bodies of 74-90 larvae from all mesocosms to detect the presence of eugregarines, *Farinocystis*, or both. We observed no accidental contamination, in the sense that all larvae from uninfected mesocosms or single-infection mesocosms were negative for non-focal parasites (**Table 1**). There was no significant difference in the prevalence of either eugregarines (single: 18.8%, coinfected: 24.3%; X^2^ = 0.71, p = 0.40) or *Farinocystis* (single: 58.4%, coinfected: 54.1%, X^2^ = 0.30, p = 0.59) among single- or co-infected mesocosms. To derive an expected prevalence of coinfection we multiplied together the prevalences from the two single-infection mesocosms (11%). The actual prevalence of coinfection was higher than expected (17.6%) but not quite significantly so (X^2^ = 3.5, p = 0.061). Within eugregarine-positive samples (**Fig. S1**), the coinfected individuals had a higher median parasite intensity than either single-infected or *Farinocystis*-negative individuals from the coinfection groups, but there was no significant difference in parasite intensity across treatment groups (Kruskal-Wallis, X^2^ = 4.6, df = 2, p = 0.10).

**Table 1.** Number and proportion of *T. castaneum* larvae infected with a focal parasite under single and coinfection conditions.

| Focal parasite | Mesocosm | Total larvae | Uninfected | Infected | Prevalence |
| --- | --- | --- | --- | --- | --- |
| *Eugregarines* | uninfected | 90 | 90 | 0 | 0 |
|  | eugregarines | 80 | 65 | 15 | 18.8 |
|  | farinocystis | 77 | 77 | 0 | 0 |
|  | coinfected | 74 | 56 | 18 | 24.3 |
| *Farinocystis* | uninfected | 90 | 90 | 0 | 0 |
|  | eugregarines | 80 | 80 | 0 | 0 |
|  | farinocystis | 77 | 32 | 45 | 58.4 |
|  | coinfected | 74 | 34 | 40 | 54.1 |
| *Both parasites* | uninfected | 90 | 90 | 0 | 0 |
|  | eugregarines | 80 | 80 | 0 | 0 |
|  | farinocystis | 77 | 77 | 0 | 0 |
|  | coinfected | 74 | 61 | 13 | 17.6 |
|  | expected | 74 | 66 | 8 | 11 |

**Notes:** the Chi Square statistics for the first two headings compare the prevalence of the focal parasite (eugregarines OR farinocystis) in single vs coinfection mesocosms, while the coinfecting heading compares the observed prevalence in the coinfecting treatment to the expected prevalence gained by multiplying parasite prevalences from the two single infection mesocosms together. Eugregarines focal: $\chi^2 = 0.71$ , $p = 0.40$ ; Farinocystis focal: $\chi^2 = 0.30$ , $p = 0.59$ ; Coinfecting expected vs observed: $\chi^2 = 3.5$ , $p = 0.061$ .

### The theoretical influence of infection on population stability over cannibalism rates

When we added infection into the LPA model and bifurcated over the cannibalism parameter (*c*_*pa*_) that controls the stability transition in disease-free populations (**Fig. 1**), we found qualitative differences in the transitions to and beyond the chaotic regime across six scenarios: infection-free, single-infection with an obligate killer, single infection a direct transmission parasite, coinfection with two obligate killers, co-infection with two directly transmitted parasites, and co-infection with one obligate killer and one directly transmitted parasite (**Figs. 1A-F**). Relative to the disease-free scenario (**Fig. 1A**), a single infection with either parasite (**Figs. 1B, C**) significantly increases stability when *c*_*pa*_ < 0.1. In contrast, co-infection produces its greatest stabilizing effect at intermediate values (0.1 < *c*_*pa*_ < 0.3) for both obligate killer and mixed transmission (**Figs. 1E, F)**, while direct transmission exhibits oscillatory behavior within the same range **(Fig. 1D**). As *c*_*pa*_ increases further, the relative stability of co-infection scenarios declines **(Figs. 1D-F**) but single-infection dynamics (**Figs. 1A-C**) are substantially more stable than coinfection at the highest parameter range.

**Figure 1.**
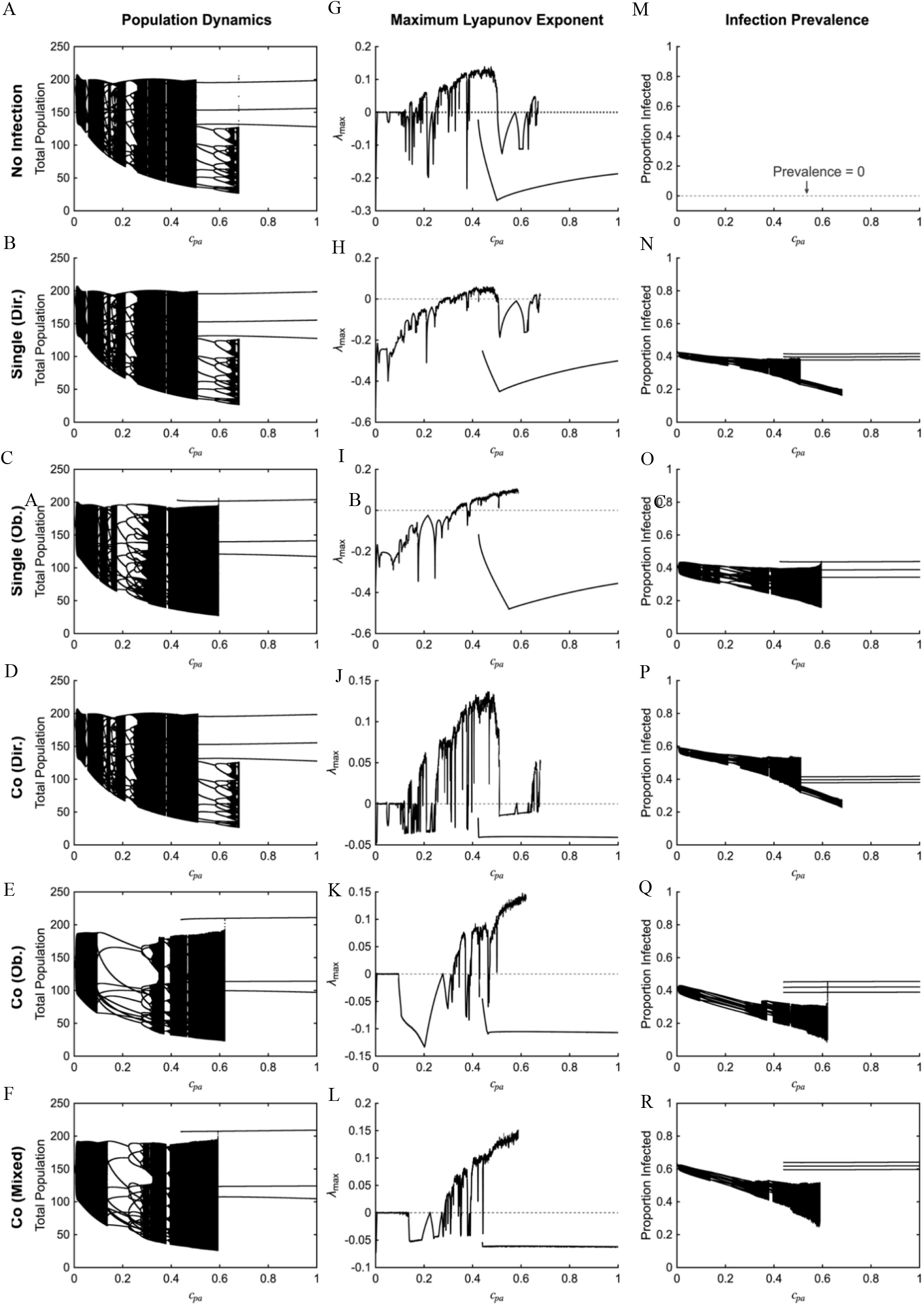
Dynamical stability across infection states. Bifurcation diagrams showing long-term total population size (Figs. 1A-F) and corresponding Lyapunov exponents (Figs. 1G-L) as a function of pupal cannibalism strength (*c*_*pa*_). The parameter *c*_*pa*_ represents density-dependent cannibalism on pupa, a key demographic interaction regulating recruitment into the adult stage and used here as the bifurcation parameter controlling population dynamics. Panels depict six infection regimes: disease-free dynamics (Figs. 1A, G, and M); single infection by a direct transmission parasite (Figs. 1B, H, and N); single infection by an obligate killer pathogen (Figs. 1C, I, and O); co-infection with direct transmission pathogens (Figs. 1D, J, and P); co-infection with obligate killer pathogens (Figs. 1E, K, and Q); and mixed transmission (Figs. 1F, L, and R). Lyapunov exponents (*λ*) quantify dynamical stability, where *λ* < 0 indicates stable equilibria, *λ* ≈ 0 marks quasiperiodicity, and *λ* >0 indicates chaotic dynamics. Differences among infection states reveals that parasite identity and transmission mode alter the onset and extent of instability. In particular, single infection with a directly transmitted parasite (Fig. 1H) shifts the stability boundary relative to single infection with an obligate killer pathogen (Fig. 1I), indicating altered sensitivity of population dynamics to cannibalism strength, whereas co-infection scenarios (Fig. 1J, K and L) produce distinct regimes not observed under single infections.

Evidence for multiple co-existing attractors is observed across all six scenarios over an intermediate range of (*c*_*pa*_) (**Figs. 1G-L**), as originally established in [33]. However, the dynamic structure of this regime differs substantially among models. In the disease-free state, as well as under single- and co-infection with direct transmission pathogens, dynamics near a *c*_*pa*_ value of 0.5 reorganize into stable periodic cycles before undergoing a second transition to chaos (**Figs. 1G, H, and J**). This contrasts with all single and co-infection dynamics involving an obligate killer, in which this re-emergence is not resolved, and the system transitions directly into chaos **(Figs. 1I, K, and L**). Overall, single infections consistently exhibited greater stability than coinfections (**Fig. S2**). For all scenarios, disease prevalence consistently mirrors population dynamics **(Figs. 1M-R**), and demonstrates that transitions among states are not simply due to the persistence or extinction of the parasites. Average parasite prevalence for all scenarios using the default parameters in **Table S1** is available in **Fig. S3**.

### The influence of transmission mode on stability

We aimed to test whether the mode of parasite transmission affects the sensitivity of population stability by bifurcating over the two parameters that ranked first and second in the global sensitivity analyses for each scenario (**Fig. S4-S7**): the focal cannibalism parameter (*c*_*pa*_) and the disease-induced mortality rate of the more virulent parasite (α_i_). The stability of populations was much more sensitive to variation in disease-induced mortality, after accounting for the effect of *c*_*pa*_, under obligate killer scenarios (**Fig. 2A, B**) relative to direct transmission **(Fig. 2C, D**). At low cannibalism intensities (*c*_*pa*_≤ 0.2) and disease-induced mortality rates (α < 0.4), both transmission modes revealed comparatively uniform dynamical profiles, spanning regions of stability and quasi periodicity across the mortality axis. As host cannibalism increased (0.2 ≤ *c*_*pa*_≤ 0.6) marked differences in sensitivity to mortality were observed across transmission modes. Obligate-killer dynamics displayed fragmented bands of stability, quasiperiodicity, and chaos, with small changes in mortality producing abrupt dynamical regime shifts. This behavior contrasted sharply with scenarios that included only directly transmitted parasites, which retained a more uniform structure. The introduction of a directly transmitted parasite into a system already infected with an obligate killer pathogen (sensitivity analysis in **Fig. 2E**) was linked to increased instability but otherwise resembled the obligate killer scenario (**Fig. 2F**), suggesting that it is the presence and virulence of the obligate killer rather than multiplicity of infection that drives stability profiles.

**Figure 2.**
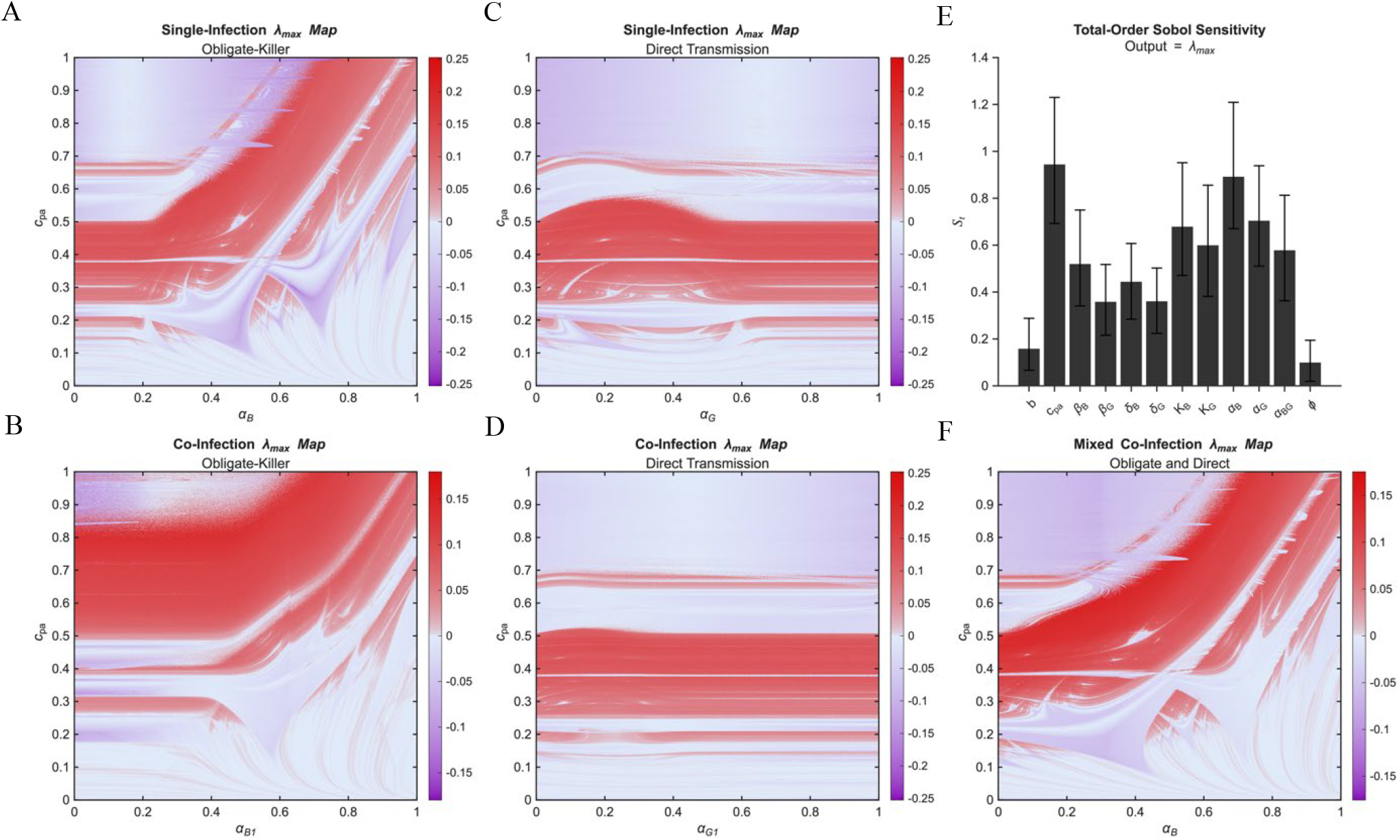
Maximum Lyapunov exponent map across cannibalism-on-pupae and disease-induced mortality rate. Single-and Co-infection dynamic regimes were classified based on their stability across different transmission modes. Color bars represent regions of chaos (red), quasi-periodicity (white), and stability (purple). Based on the results from global sensitivity analysis, two focal parameter pairs were chosen for each model due to their importance for population stability (Figure S4-S7). Models were bifurcated across cannibalism intensity (*c*_*pa*_) and disease-induced mortality (*α*). Stability was more sensitive to variation in *α* than *c*_*pa*_, particularly under obligate killer conditions (A, B) compared to direct transmission (C, D). As *c*_*pa*_ increased, obligate-killer systems exhibited fragmented transitions among dynamical states, whereas direct transmission retained a more uniformly chaotic structure. Accordingly, overall dynamics in the mixed model (E, F) remained primarily driven by the virulent obligate killer.

### Interactions among parasites influences host population stability

We hypothesized that antagonistic interactions would promote stability, whereas facilitative interactions would destabilize dynamics, particularly when parasite interactions affected both spore deposition and spore infectiousness (*ϕ*_*ij*_ *β*_*i*_*E*_*i* (_*t*_)_) (See Supplementary Materials). To explore the role of parasite interactions on population dynamics, we used the default parameters (**Table S1**) to generate bifurcation diagrams for co-infected populations with two obligate killers (**Fig. 3A-C**) and under mixed transmission (**Fig. 3B-F**), which were particularly interesting based on the results from previous analyses (**Figs. 1-2)**. In the original asymmetric interaction formulation, total-order Sobol sensitivity analysis revealed that the coefficients of facilitation and antagonism (*ϕ*) had minimal impact on population stability across all models, as highlighted in the mixed transmission model from **Fig. 2E**. When we allowed parasite interactions to affect transmission parameters, total-order Sobol indices show that the antagonism-facilitation coefficient dominated the Lyapunov exponent estimation in the obligate killer model (**Fig. 3C**) but still remained modest in the mixed-transmission model (**Fig. 3F**).

**Figure 3.**
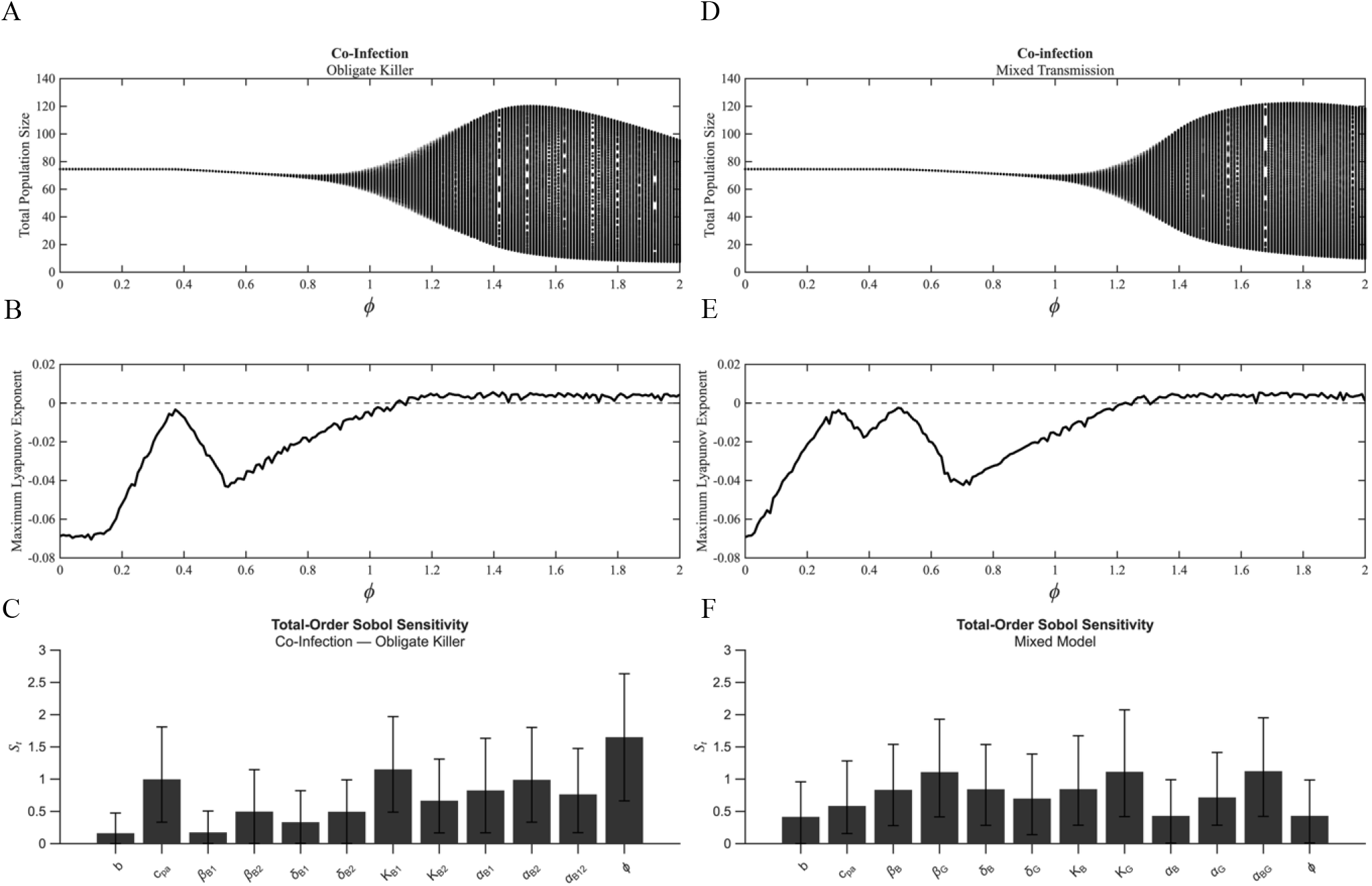
Dynamical stability profile across antagonism and facilitation. Bifurcation analysis and maximum Lyapunov exponent estimation as a function of the interaction coefficient (*ϕ*), where *ϕ*> 1 denotes facilitation, *ϕ*= 1 indicates neutral interactions, and *ϕ*< 1 represents antagonism. Panels A-D classify the co-infection dynamics of two genetically distinct obligate killer strains across varying population sizes and divergence rates for both obligate killer and mixed-transmission models. Sobol sensitivity analysis identifies the antagonism-facilitation coefficient as the primary driver of dynamical instability in the obligate killer model (E), but only a minor contributor in determining the stability of the mixed transmission model (F).

At extreme levels of antagonism (*ϕ*≤ 0.5), regardless of transmission mode, population trajectories remain tightly clustered and the maximum Lyapunov exponent is strongly negative, indicating stable long-term dynamics with little variability (**Fig. 3A-B, 3D-E**). As *ϕ*increases towards neutral interactions, the stable equilibrium begins to lose robustness. Population trajectories gradually broaden, and equilibrium values shift towards quasiperiodicity and mild chaos. This transition is observed in the rise of the λ_max_ toward zero, signaling a reduction in local stability (**Figs. 3B, E**). Beyond *ϕ*= 1 (neutral interactions giving way to facilitation), total population dynamics change qualitatively for both scenarios; the maximum Lyapunov exponent fluctuates around zero and becomes intermittently positive, consistent with transitions to chaotic dynamics when parasites facilitate each other. Notably, however, the mixed-transmission model requires a higher facilitation threshold than the obligate-killer model to trigger chaotic dynamics within the system.

## Discussion

In this study, we investigated the impact of multiple infections on the stability of host population dynamics. Starting with the classic LPA model, which is both intimately grounded in *Tribolium* flour beetle biology and well-explored analytically, we performed bifurcation and sensitivity analyses to systematically characterize population stability over a range of (co)infection scenarios, paying particular attention to the role of transmission mode and facilitative and antagonistic interactions among parasites. Our model predicts that coinfection with common parasites can alter the range of demographic parameters in which the system transitions from stability to chaos, and that chaos is a common feature of outcomes when obligate killer parasites facilitate each other even in regions of parameter space that are otherwise stable in the absence of infection. Given the diversity and tractability of parasites with direct and obligate killer transmission modes in the *Tribolium* system, as demonstrated in the experiments here and other recent work [38], this study sets up concrete and testable predictions for future experimental mesocosms and highlights the general importance of knowing which demographic and infection parameters are affected by parasite-parasite interactions in any coinfection system.

A long tradition of integrated theory and experiments has solidified the expectation that parasites regulate the stability and periodicity of host population dynamics [1,2]. Theory on forest insect population dynamics, for example, emphasized that spore-forming parasites that induce rapid mortality in their hosts and decay slowly in the environment are particularly primed to induce population cycles relative to single stable equilibria. Adding an element of community complexity, a model of host population stability in the presence of two parasitoids and one parasite suggest that the infection promotes the coexistence of the parasitoids while also opening a wide range of parameter space where host population dynamics are inherently unstable or even chaotic [49]. Similarly, theoretical models of microparasite-macroparasite co-infection have shown that parasite-mediated feedbacks between transmission and host demography can fundamentally alter population stability, generating oscillatory dynamics and shifting equilibrium outcomes across parameter space [48]. In this study, we started by examining the interaction of single infections with the rate of cannibalism of adults on pupae; in the original disease-free model, this parameter exhibited a dynamic series of transitions from stability to quasi-periodicity to chaos [33,34,50]. We found that single infection with either directly transmitted or obligate killer parasites provided a stabilizing influence on population dynamics when the cannibalism rate was both low and high. However, it exacerbated instability at intermediate cannibalism values, suggesting the importance of density-dependent larval abundance in regulating feedback loops and stage transitions but also indicates a localized dynamical transition rather than a monotonic, gradual destabilization.

When we added the obligate killer to promote multiple infections within the population, we observed an increase in stability at low to intermediate cannibalism rates relative to either disease-free or single-infection scenarios, characterized mainly by a greater range of more regular population cycles. This contrasted with the direct transmission profile, where single infection was more stable across the designated *c*_*pa*_ parameter space (**Fig. S2**). Interestingly, the double-bifurcation analyses on parasite-induced mortality rate versus cannibalism rate suggested that transmission mode strongly interacts with the multiplicity of infection. Changes in the disease-induced mortality rates of directly transmitted parasites barely altered the stability landscape, with most of the variance attributable to cannibalism rate. Co-infection with mixed pathogens, on the other hand, presented a similar profile to obligate killer single infections, which shift the region of chaos into higher cannibalism rates when disease-induced mortality (α) was low and interacted with cannibalism as α increased further. This opened up a region of stability in the parameter space where cannibalism rate is realistically low (*c*_*pa*_<0.2) and disease-induced mortality is relatively high (**Fig. 2B**,**F**). Notably, many obligate-killer infections exhibit parameter ranges that fall within this region; previous estimates for baculovirus-induced mortality in moths range from 0.4-0.8/week (about 10-fold higher than background mortality rates) and mortality rates from *Bacillus thuringiensis* are even higher than that [2]. While we did not comprehensively estimate mortality rates from *Farinocystis tribolii* in this study, anecdotally we observed that it wipes out the majority of infected larvae and is likely on par with baculovirus mortality rates. Thus, while coinfection exerts complex effects on stability when scanning across the entire parameter space, we predict that in natural insect populations coinfection will have the net effect of promoting population stability.

The exception, of course, is if one parasite facilitates another, particularly through promoting the infectiousness of transmission-stage spores. When we allowed the interaction coefficient (*ϕ*) to promote or antagonize spore infectivity (*ϕ*_*ij*_ *β*_*i*_*E*_*i*(_*t*_)_) as well as spore production (e.g. (*ϕ*_*ij*_*α*_*ij*_*L*_*ij*_(*t*)) for obligate killer parasites, we found that facilitation (*ϕ*> 1) promotes a transition into unpredictable population cycles. Interestingly, it takes barely any facilitation to promote chaos under two obligate killers, while mixed transmission mode coinfections require substantially higher facilitation rates. There are very few coinfection studies from any model system that employ such a lethal combination as most focus on mixed transmission modes or two directly transmitted parasites. In the case of eugregarines and *Farinocystis*, our data suggest that the two parasites may weakly facilitate each other, given that rates of co-infection are higher than expected and that *Farinocystis*-coinfected parasites have higher median eugregarine loads, although neither effect reached significance and the effect size is likely not high enough to promote chaos. Previous work with eugregarines and a different obligate killer, *Bacillus thuringiensis*, demonstrated that coinfection with eugregarines promoted a 39% higher rate of infection-induced mortality when *Tribolium* beetles were infected with the bacterium [38]. The number of bacteria during the acute phase of infection was not significantly different between Bt-only and co-infected groups, however, suggesting that the mortality effect reflected a reduction in tolerance [38] without a concomitant increase in bacterial load. While that study did not measure changes in spore production and infectivity, our study highlights that these parameters would directly affect predictions for stability under coinfection and should be addressed in future work.

We did make several assumptions in this study that should be carefully considered, however. Most notably, the discrete-time structure provides less flexibility to modify larval development times; the most obvious phenotypic evidence of parasitism from eugregarines is a modest extension of development time [38], particularly under low-resource conditions, which would otherwise affect the buildup of larvae in the population. While our model explicitly captures the life cycle of beetles, it may be less applicable to systems with overlapping generations, such as those typical of most mammals. Conversely, it may extend readily to other holometabolous organisms that share comparable stage-structured life cycles to those of our focal system.

Taken together, our results demonstrate that co-infection can fundamentally reshape population stability, generating outcomes that span from increased stability to attenuated chaos in stage-structured populations. Rather than acting solely as an additional source of mortality, co-infection emerges as a dynamical perturbation whose effects depend on how pathogens transmit and interact within hosts. By connecting these mechanisms to emergent stability patterns, our framework provides a mechanistic explanation for why otherwise similar host-pathogen systems can display contrasting population dynamics. Explicitly parametrizing these processes using experimentally derived estimates will be essential for translating theoretical predictions into quantitative forecasts of population behavior. Such integration may ultimately improve our ability to anticipate disease-driven fluctuations relevant to conservation, biological control, and management of naturally variable populations.

## Acknowledgements

We thank Robert Desharnais for providing MATLAB code from the LPA model and Faith Rovenolt for preliminary exploration of the model. This study was funded by NSF DEB award 2508355 to A.T.T.

## List of Supplementary Materials

### Supplemental Tables

**Table S1**. Parameter values used in the LPA and infection-extended models

**Table S2**. Sobol parameter ranges defined over the full set of models

### Supplemental Figures

**Fig.S1**. Intensity of eugregarine infection in larvae

**Fig.S2**. Differential stability across transmission mode and type of infection

**Fig.S3**. Prevalence plot across infection domains

**Fig.S4**. Global sensitivity analysis for single infection under obligate killer transmission

**Fig.S5**. Global sensitivity analysis for single infection under direct transmission

**Fig.S6**. Global sensitivity analysis for obligate-killer co-infection

**Fig.S7**. Global sensitivity analysis for co-infection involving two direct transmission pathogens

## Notes

### Competing Interest Statement

The authors have declared no competing interest.

