## Supplemental Material for "The impact of coinfection on population stability and chaos"

#### *Supplementary methods for analyzing prevalence*

Simulations were run for 1000 time-steps, discarding the first 100 to remove transient dynamics. Prevalence was calculated as the proportion of infectious individuals relative to the total larval population. The calculation was performed across six scenarios: (i) Disease-free infection, (ii) Single infection with direct transmission, (iii) Single infection with obligate killer transmission, (iv) co-infection with direct transmission, (v) co-infection with obligate-killer transmission, and (vi) mixed model co-infection. All simulations used a cannibalism-on-pupa value that approximated the realistic $c_{pa}$ of 0.0047 (Table $S1$).

#### *Supplementary methods for analyzing Sobol Sensitivity*

For each of the four models, we quantified global parameter sensitivity using variance-based Sobol indices with the maximum Lyapunov exponent as the response variable [19]. In each case, parameters were assigned an independent uniform distribution over prespecified biologically plausible bounds. Fixed host demographic parameters were kept constant during the runs. To efficiently sample the parameter space, we used quasi-Monte Carlo Sobol sequence with skip and leap setting and Matoušek–Affine–Owen scrambling to reduce correlation artifacts. For each sequence, we generated 2N Sobol sequence points, split into two independent sampling matrices A and B, and linearly mapped each column to the corresponding parameter bounds. Following Saltelli design, we computed the maximum Lyapunov exponent from the corresponding map. Simulations were initiated with the burn-in steps, after which the maximum Lyapunov exponent was estimated, using a finite-difference tangent-vector method. At each step, the map at the current and perturbed state were estimated. The perturbation direction was renormalized, the logarithm of the state vector was accumulated and averaged over time.

Sobol first-order and total-order were computed using Saltelli estimators. For each parameter, $S_I$ quantified the fraction of variance in maximum Lyapunov exponent attributable to a parameter on its own, while $S_T$ quantified the fraction attributable to a certain parameter, including all interaction effects with other parameters. To quantify uncertainty and showcase error bars, we used a nonparametric bootstrap over the base samples. All analyses showcased error bars and were performed in MATLAB.

#### *Model limitations and considerations*

For the model to be successfully applied across different systems and ensure broad applicability, it is important to consider biological variability, such as differences in parasite prevalence within already characterized host-pathogen systems. For instance, in the current study, eugregarine prevalence was 20%, while previous studies have reported it as high as 60% [Schulz 2025]. To accommodate this variability and improve the model's generalizability, we aimed for an average prevalence of 40%, which balances these extremes (Fig. S3).

### Supplementary Tables

**Table S1. Parameter values used in the LPA and infection-extended models**. Parameter definitions, baseline values, and literature sources used for all simulations. Parameter values lacking experimental estimates were estimated by calibration to reproduce infection prevalence consistent with experimental observations (Simpson, unpublished data).

| State variable | Definition | Value | Reference |
|---|---|---|---|
| $L_{tot(t)}$ | Total population at time *t* (Sum of all larval classes in that model) | 250 | (Constantino et al., 1997) |
| $L_{s(t)}$ | Susceptible larvae at time *t* | 240 | (Constantino et al., 1997) |
| $L_{i(t)}$ or $L_{j(t)}$ | Infected larvae with *Bt* at time *t* | 10 | (Constantino et al., 1997) |
| $L_{i1(t)}$ or $L_{j1(t)}$ | Initial larvae infected with strain one (co-infection) | 5 | Estimated |
| $L_{i2(t)}$ or $L_{j2(t)}$ | Initial larvae infected with strain two (co-infection) | 4 | Estimated |
| $L_{i1i2(t)}$ or $L_{j1j1(t)}$ | Initial co-infected larval population | 1 | Estimated |
| $E_{i1(t)}$ or $E_{j1(t)}$ | Initial environmental contamination (strain one) | 0.2 | Estimated |
| $E_{i2(t)}$ or $E_{j2(t)}$ | Initial environmental contamination (strain two) | 0.2 | Estimated |
| $P_t$ | Number of pupae at time *t* | 5 | (Constantino et al., 1997) |
| $A_t$ | Number of adults at time *t* | 100 | (Constantino et al., 1997) |
| $b$ | Number of larval recruits per adult per unit of time in the absence of cannibalism | 6.598 | (Constantino et al., 1997) |
| $c_{el}$ | Cannibalism of eggs by larvae | 0.01209 | (Constantino et al., 1997) |
| $c_{ea}$ | Cannibalism of eggs by adults | 0.01155 | (Constantino et al., 1997) |
| $c_{pa}$ | Cannibalism of pupae by adults | 0.0047 | (Constantino et al., 1997) |
| $\mu_l$ | Death rate of larvae | 0.2055 | (Constantino et al., 1997) |
| $\mu_a$ | Death rate of adults | Naturally: 0.007629 Experimentally set to: 0.96 | (Constantino et al., 1997) |
| $\phi_{k1}$and $\phi_{k2}$ | General antagonism-facilitation coefficient | $\phi \in$ 0-2 | Estimated |

**Table S2. Sobol parameter ranges defined over the full set of models**. List of parameter ranges evaluated for sensitivity analysis. The presence of multiple plausible parameter combinations implies that system parameter may significantly vary across parameter space, highlighting the need for experimental parameterization to better constrain predictions.

| Parameter | Range |
|---|---|
| $b$ | 5.5 - 7.5 |
| $c_{pa}$ | 0 - 1 |
| $\alpha_k$ and $\alpha_{k1k2}$ | 0 - 1 |
| $\delta_i$ | 0.02 – 0.10 |
| $\delta_j$ | 0.02 – 0.15 |
| $\kappa_i$ | 0.001 – 0.03 |
| $\kappa_j$ | 0.001 – 0.010 |
| $\beta_i$ | 0.90 – 1.30 |
| $\beta_j$ | 0.80 – 1.30 |
| $\phi_{k1}$ and $\phi_{k2}$ | 0 - 2 |

### Supplementary Figures

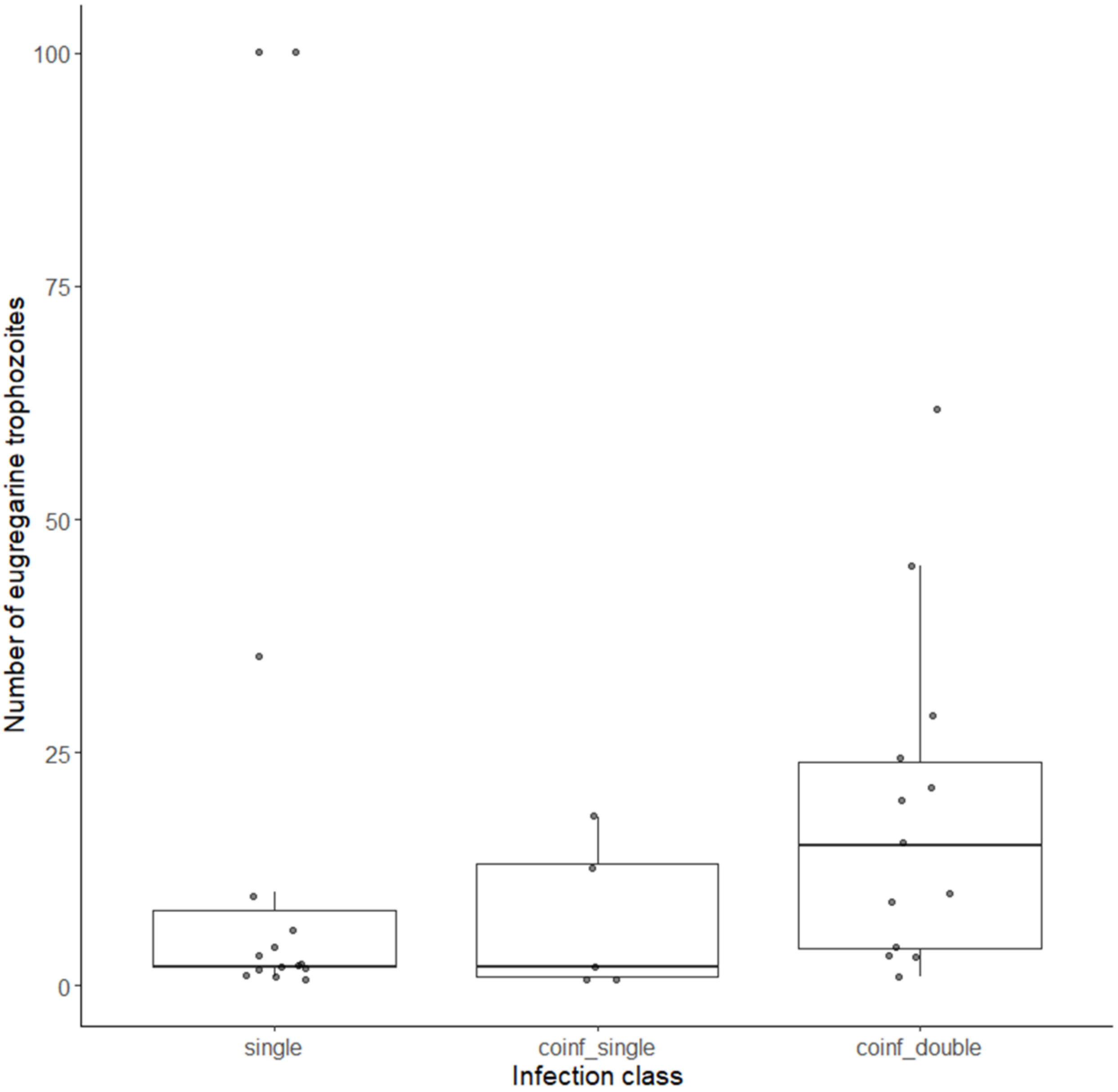


**Fig.S1. Intensity of eugregarine infection in larvae.** The number of visible eugregarine trophozoites was counted by microscopy for each individual that had at least one trophozoite from each of the infection mesocosms. There were no significant differences among groups (Kruskal-Wallis, $X^2 = 4.6$, df = 2, p = 0.10).

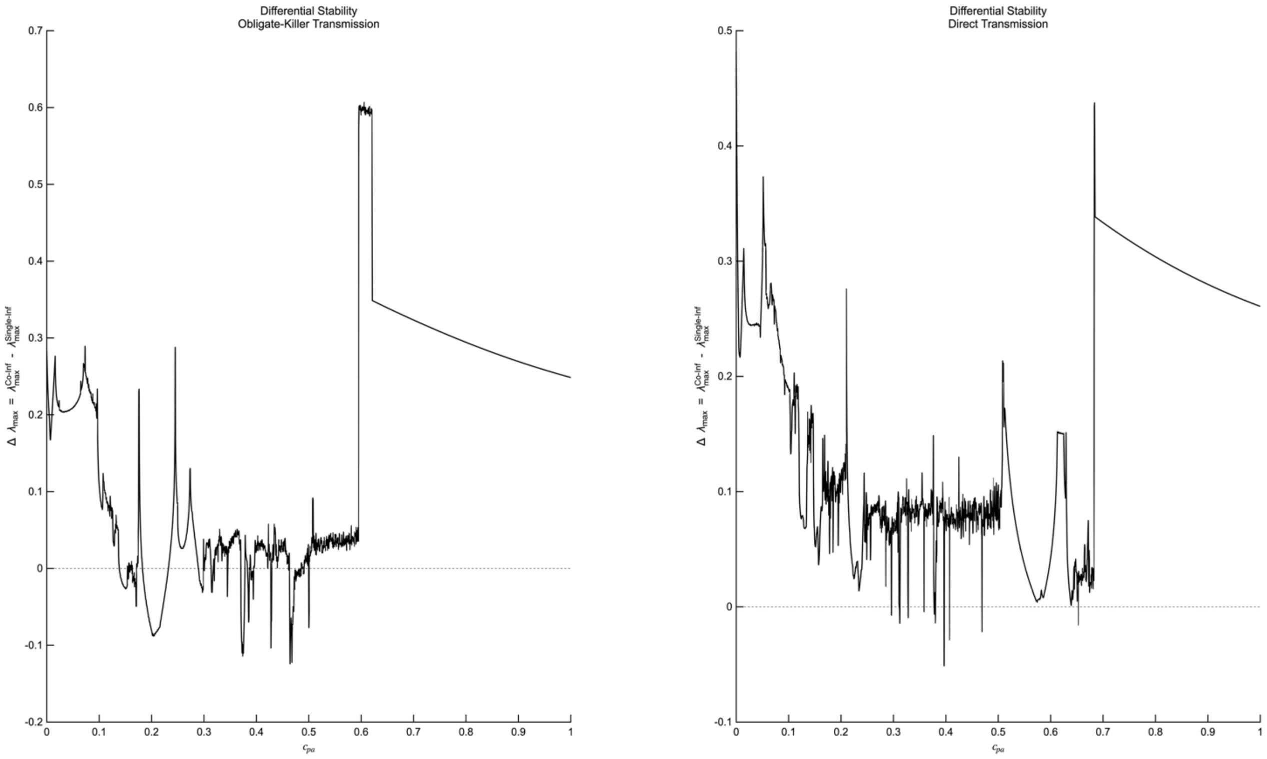


**Fig.S2. Differential stability across transmission mode and type of infection.** Stability differences between single- and co-infection across transmission mode reveal single exposure as unanimously more stable. Using forward sweeps from the simulations shown in figure one, differential stability was calculated by subtracting the maximum Lyapunov exponent of the single-infection from that of the co-infection model across each transmission type. Negative values indicate regions where co-infection is more stable, whereas positive values indicate regions where single infections are more stable.

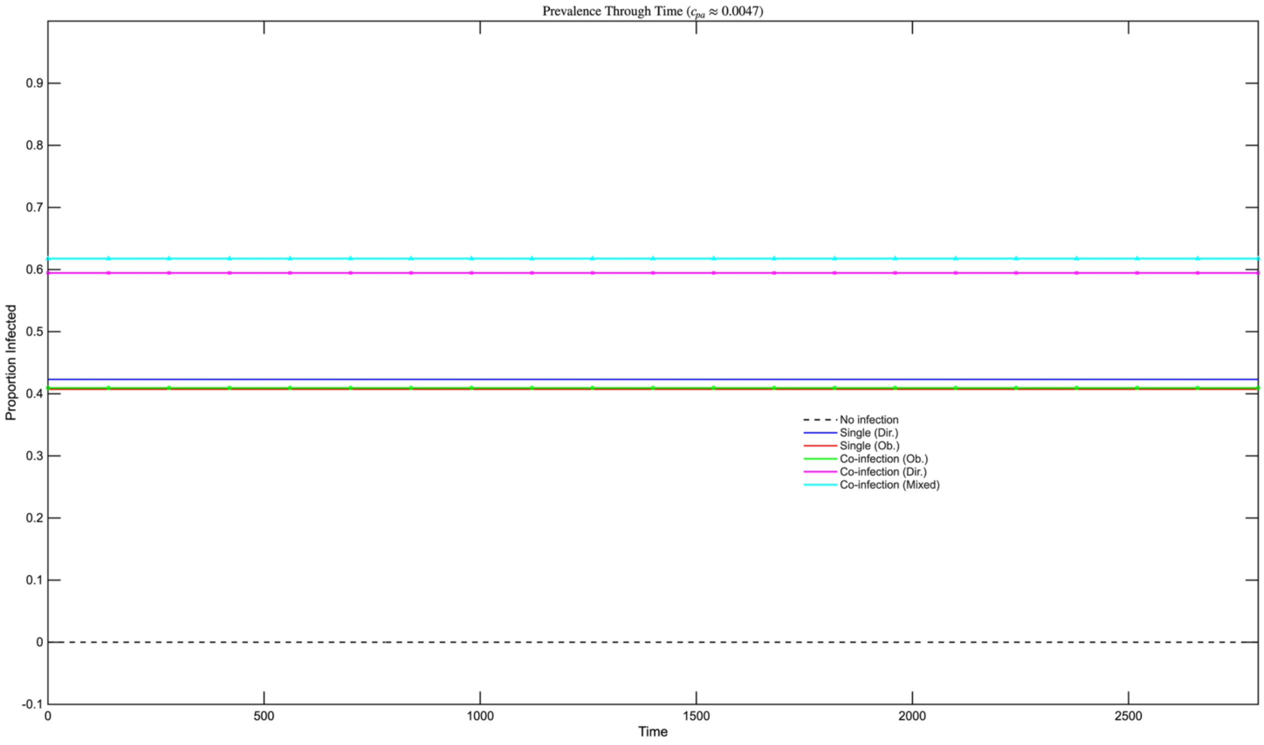


**Fig.S3. Prevalence plot across infection domains.** Infection prevalence was quantified as proportion of infected larvae relative to total larval abundance through time using the parameters listed in Supplementary Table S2. Prevalence dynamics were compared across six epidemiological scenarios: disease-free, single infection with an obligate killer, single infection with a direct transmission parasite, co-infection with two obligate killers, co-infection with two direct transmission parasites, and mixed-mode co-infection at the parametrized $c_{pa}$ value of 0.0047.

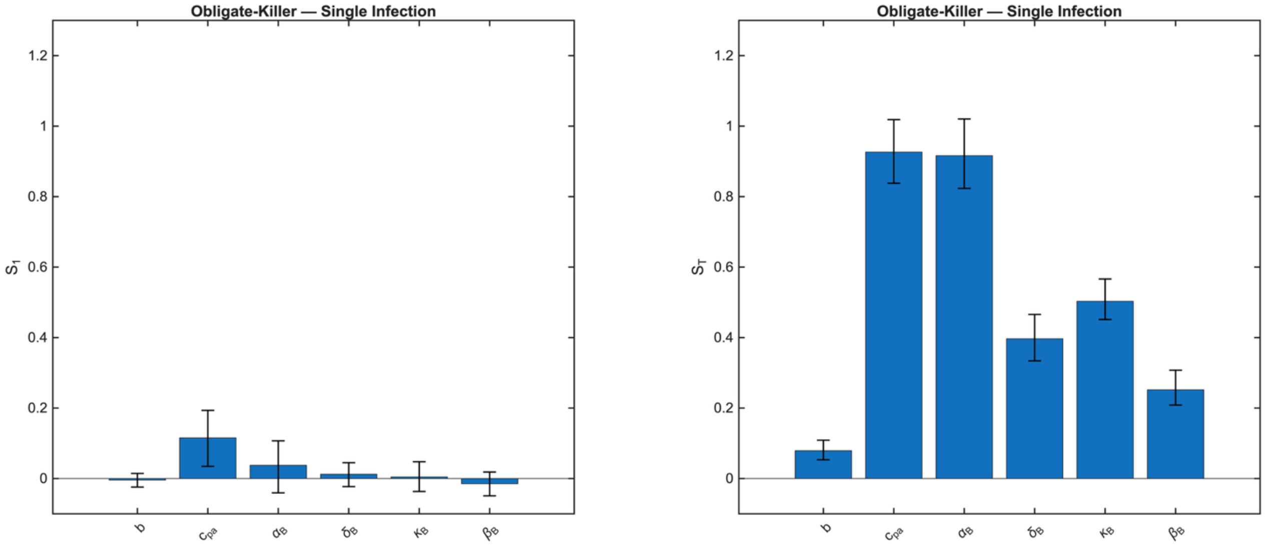


**Fig.S4. Global sensitivity analysis for single infection under obligate killer transmission.** First-order Sobol sensitivity analysis in single infected obligate-killer beetles revealed that each parameter alone had a modest effect on the maximum Lyapunov exponent, while cannibalism-on-pupa and the single-infection-induced mortality rate were predominant when interactions with other parameters were included.

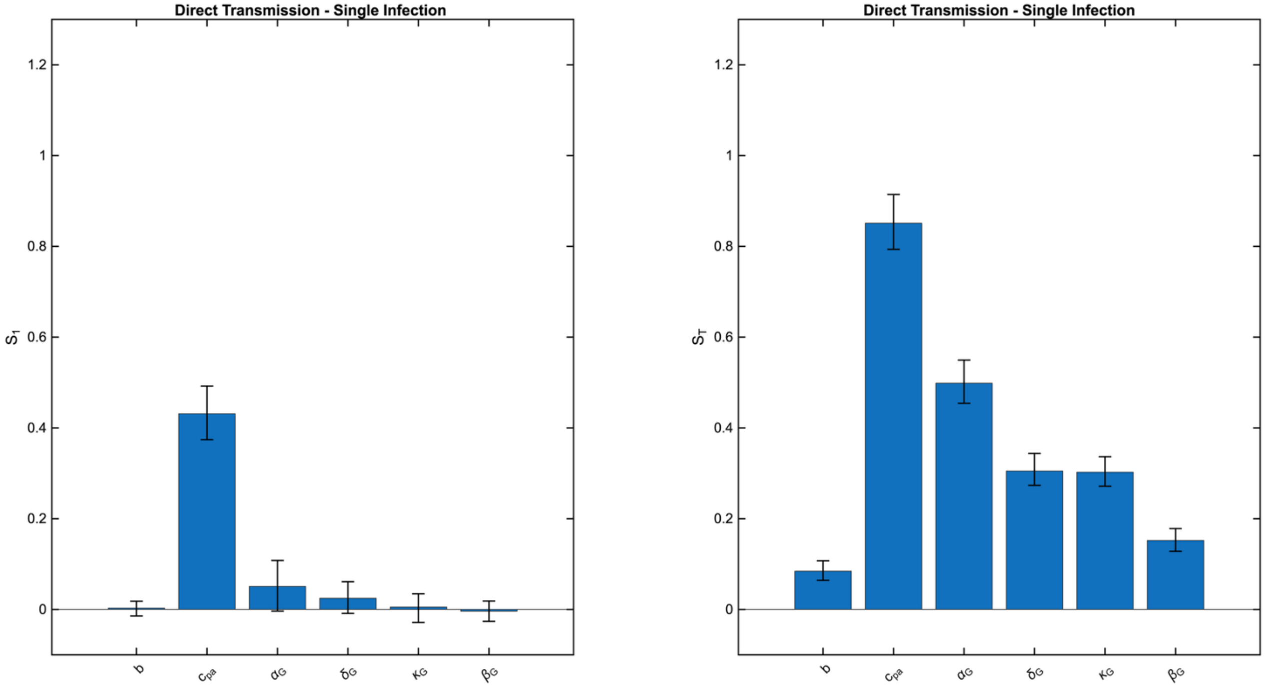


**Fig.S5. Global sensitivity analysis for single infection under direct transmission.** First-order Sobol sensitivity analysis revealed a strong influence of cannibalism-on-pupa both on its own and in combination with other sampled parameters. The single infection-induced mortality rate modestly influenced the maximum Lyapunov exponent when interacting with other variables.

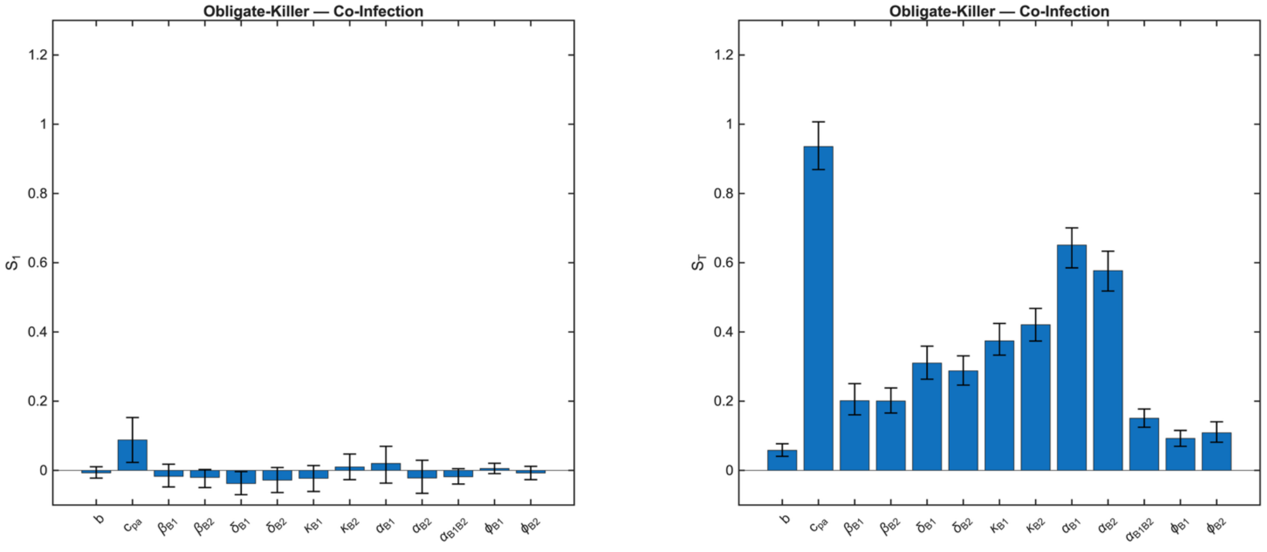


**Fig.S6. Global sensitivity analysis for two obligate-killer co-infection**. First-order Sobol sensitivity analysis revealed minimal influence of each sampled parameter. Cannibalism-on-pupa and the single infection-induced mortality rate were predominant when interactions with other parameters were included.

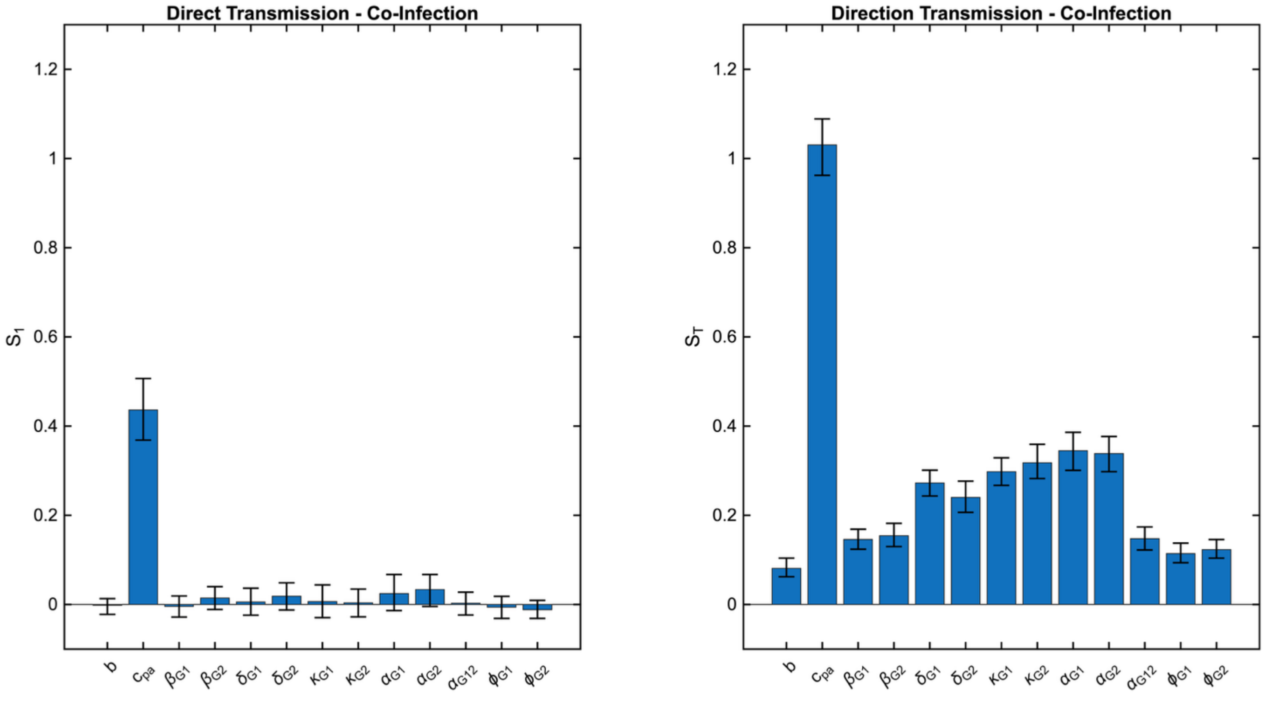


**Fig.S7. Global sensitivity analysis for co-infection involving two direct transmission pathogens.** First- and total-order Sobol indices revealed a strong influence of cannibalism-on-pupa and a modest influence of the single infection-induced mortality rate.

**Supplementary Equations: Co-infection model formulation**

### Homologous coinfecting parasites (obligate killer *or* direct transmission)

$$L_{tot(t)} = L_{S(t)} + L_{k_1(t)} + L_{k_2(t)} + L_{k_1k_2(t)} \quad \text{Eq. S1}$$

$$R_t = bA_t e^{-C_{el}L_{tot(t)} - C_{ea}A_t} \quad \text{Eq. S2}$$

$$L_{S(t+1)} = R_t e^{-(\beta_{k_1}E_{k_2(t)} + \beta_{k_2}E_{k_2(t)})} \quad \text{Eq. S3}$$

$$L_{I_1(t+1)} = R_t\left(1 - e^{-\beta_{k_1}E_{k_1(t)}}\right)e^{-\beta_{k_2}E_{k_2(t)}} \quad \text{Eq. S4}$$

$$L_{I_2(t+1)} = R_t\left(1 - e^{-\beta_{k_2}E_{k_2(t)}}\right) \quad \text{Eq. S5}$$

$$L_{I_1I_2(t+1)} = R_t\left(1 - e^{-\beta_{k_1}E_{k_1(t)}}\right)\left(1 - e^{-\beta_{k_2}E_{k_2(t)}}\right) \quad \text{Eq. S6}$$

$$P_{t+1} = (1 - \mu_l)L_{S(t)} + \left(1 - \mu_l - \alpha_{k_1}\right)L_{k_1(t)} + \left(1 - \mu_l - \alpha_{k_2}\right)L_{k_2(t)} + \left(1 - \mu_l - \alpha_{k_1k_2}\right)L_{k_1k_2(t)} \quad \text{Eq. S7}$$

$$A_{t+1} = P_t e^{-C_{pa}A_t} + A_t(1 - \mu_a) \quad \text{Eq. S8}$$

where $k \in \{i, j\}$ denotes parasite identity ($i$, obligate killer; $j$, direct transmission)

### Obligate Killer

$$E_{i1(t+1)} = (1 - \delta_{i1})E_{i1(t)} + \left(1 - E_{i1(t)}\right)\left(1 - e^{-K_{i1}\left(\alpha_{i1}L_{i1(t)} + \phi\alpha_{i1i2}L_{i1i2(t)}\right)}\right) \quad \text{Eq. S9a}$$

$$E_{i2(t+1)} = (1 - \delta_{i2})E_{i2(t)} + \left(1 - E_{i2(t)}\right)\left(1 - e^{-K_{i2}\left(\alpha_{i2}L_{i2(t)} + \phi\alpha_{i1i2}L_{i1i2(t)}\right)}\right) \quad \text{Eq. S10a}$$

### Direct Transmission

$$E_{j1(t+1)} = \left(1 - \delta_{j1}\right)E_{j1(t)} + \left(1 - E_{j1(t)}\right)(1 - e^{-\mathrm{K}_{j1}\left[(1-\alpha_{j1})L_{j1(t)} + \phi_{k1}(1-\alpha_{j1j2})L_{j1j2}(t)\right]}) \quad \text{Eq. S9b}$$

$$E_{j2(t+1)} = \left(1 - \delta_{j2}\right)E_{j2(t)} + \left(1 - E_{j2(t)}\right)(1 - e^{-\mathrm{K}_{j2}\left[(1-\alpha_{j2})L_{j2(t)} + \phi_{k2}(1-\alpha_{j1j2})L_{j1j2}(t)\right]}) \quad \text{Eq. S10b}$$

### Symmetric Interaction Formulation:

$$\phi_k\beta_k E_{k(t)}$$

For the analysis of antagonism and facilitation in figure 3, a symmetric interaction model was formulated. All instances of the transmission term $\beta_k E_{k(t)}$ were replaced with the symmetric interaction form $\phi_k\beta_k E_{k(t)}$ as shown below:

$L_{tot(t)} = L_{S(t)} + L_{k_1(t)} + L_{k_2(t)} + L_{k_1k_2(t)}$ Eq. S11

$R_t = bA_t e^{-C_{el}L_{tot(t)} - C_{ea}A_t}$ Eq. S12

$L_{S(t+1)} = R_t e^{-(\phi_k \beta_{k_1} E_{k_2(t)} + \phi_k \beta_{k_2} E_{k_2(t)})}$ Eq. S13

$L_{I_1(t+1)} = R_t\left(1 - e^{-\phi_k \beta_{k_1} E_{k_1(t)}}\right) e^{-\phi_k \beta_{k_2} E_{k_2(t)}}$ Eq. S14

$L_{I_2(t+1)} = R_t\left(1 - e^{-\phi_k \beta_{k_2} E_{k_2(t)}}\right)$ Eq. S15

$L_{I_1 I_2(t+1)} = R_t\left(1 - e^{-\phi_k \beta_{k_1} E_{k_1(t)}}\right)\left(1 - e^{-\phi_k \beta_{k_2} E_{k_2(t)}}\right)$ Eq. S16

$P_{t+1} = (1 - \mu_l) L_{S(t)} + \left(1 - \mu_l - \alpha_{k_1}\right) L_{k_1(t)} + \left(1 - \mu_l - \alpha_{k_2}\right) L_{k_2(t)} + \left(1 - \mu_l - \alpha_{k_1 k_2}\right) L_{k_1 k_2(t)}$ Eq. S17

$A_{t+1} = P_t e^{-C_{pa}A_t} + A_t(1 - \mu_a)$ Eq. S18

where $k \in \{i, j\}$ denotes parasite identity ($i$, obligate killer; $j$, direct transmission)

### Obligate Killer

$E_{i1(t+1)} = (1 - \delta_{i1}) E_{i1(t)} + \left(1 - E_{i1(t)}\right)\left(1 - e^{-K_{i1}\left(\alpha_{i1} L_{i1(t)} + \phi_k \alpha_{i1i2} L_{i1i2(t)}\right)}\right)$ Eq. S19a

$E_{i2(t+1)} = (1 - \delta_{i2}) E_{i2(t)} + \left(1 - E_{i2(t)}\right)\left(1 - e^{-K_{i2}\left(\alpha_{i2} L_{i2(t)} + \phi_k \alpha_{i1i2} L_{i1i2(t)}\right)}\right)$ Eq. S20a

### Direct Transmission

$E_{j1(t+1)} = \left(1 - \delta_{j1}\right) E_{j1(t)} + \left(1 - E_{j1(t)}\right)\left(1 - e^{-\mathrm{K}_{j1}\left[(1-\alpha_{j1}) L_{j1(t)} + \phi_{k1}(1-\alpha_{j1j2}) L_{j1j2}(t)\right]}\right)$ Eq. S19b

$E_{j2(t+1)} = \left(1 - \delta_{j2}\right) E_{j2(t)} + \left(1 - E_{j2(t)}\right)\left(1 - e^{-\mathrm{K}_{j2}\left[(1-\alpha_{j2}) L_{j2(t)} + \phi_{k2}(1-\alpha_{j1j2}) L_{j1j2}(t)\right]}\right)$ Eq. S20b

***Mixed transmission mode model formulation (Obligate killer + Direct transmission)***

$L_{tot(t)} = L_{S(t)} + L_{i(t)} + L_{j(t)} + L_{ij(t)}$ Eq. S21

$R_t = bA_t e^{-C_{el}L_{tot(t)} - C_{ea}A_t}$ Eq. S22

$L_{S(t+1)} = R_t e^{-(\beta_i E_{i(t)} + \beta_j E_{j(t)})}$ Eq. S23

$L_{i(t+1)} = R_t\left(1 - e^{-\beta_i E_{i(t)}}\right) e^{-\beta_j E_{j(t)}}$ Eq. S24

$$L_{j(t+1)} = R_t\left(1 - e^{-\beta_j E_{j(t)}}\right)e^{-\beta_i E_{i(t)}}$$ Eq. S25

$$L_{ij(t+1)} = R_t\left(1 - e^{-\beta_i E_{i(t)}}\right)\left(1 - e^{-\beta_j E_{j(t)}}\right)$$ Eq. S26

$$P_{t+1} = (1 - \mu_l)L_{S(t)} + (1 - \mu_l - \alpha_i)L_{i(t)} + \left(1 - \mu_l - \alpha_j\right)L_{j(t)} + \left(1 - \mu_l - \alpha_{ij}\right)L_{ij(t)}$$ Eq. S27

$$A_{t+1} = P_t e^{-C_{pa}A_t} + A_t(1 - \mu_a)$$ Eq. S28

$$E_{OB(t+1)} = (1 - \delta_{OB})E_{OB(t)} + \left(1 - E_{OB(t)}\right)\left(1 - e^{-K_{OB}\left(\alpha_{OB}L_{OB(t)} + \phi_{OB}\alpha_{DOB}L_{DOB(t)}\right)}\right)$$ Eq. S29

$$E_{D(t+1)} = (1 - \delta_D)E_{D(t)} + \left(1 - E_{D(t)}\right)\left(1 - e^{-K_D\left((1-\alpha_j)L_{j(t)} + \phi_D(1-\alpha_{DOB})L_{DOB(t)}\right)}\right)$$ Eq. S30
